# Can individual and integrated water, sanitation, and handwashing interventions reduce fecal contamination in the household environment? Evidence from the WASH Benefits cluster-randomized trial in rural Kenya

**DOI:** 10.1101/731992

**Authors:** Amy J. Pickering, Jenna Swarthout, MaryAnne Mureithi, John Mboya, Benjamin F. Arnold, Marlene Wolfe, Holly N. Dentz, Audrie Lin, Charles D. Arnold, Gouthami Rao, Christine P. Stewart, Pavani K. Ram, Thomas Clasen, John M. Colford, Clair Null

**Affiliations:** Civil and Environmental Engineering, Tufts University, Medford, MA; Innovations for Poverty Action, Nairobi, Kenya; University of California, Berkeley, Berkeley, CA; University of California, Davis, Davis, CA; University at Buffalo, Buffalo, NY; Emory University, Atlanta, GA; Mathematica Policy Institute, Washington D.C

## Abstract

Combined water, sanitation, and handwashing (WSH) interventions have the potential to reduce fecal pathogens along more transmission pathways than single interventions alone. We measured *Escherichia coli* levels in 3909 drinking water samples, 2691 child hand rinses, and 2422 toy ball rinses collected from households enrolled in a two-year cluster-randomized controlled trial evaluating single and combined WSH interventions. Water treatment alone reduced *E. coli* in drinking water, while a combined WSH intervention improved water quality by the same magnitude but did not affect levels of fecal indicator bacteria on child hands or toy balls. The failure of the WSH interventions to reduce *E. coli* along important child exposure pathways is consistent with the lack of a protective effect from the interventions on child diarrhea or child growth during the trial. Our results have important implications for WSH program design; the sanitation and handwashing interventions implemented in this trial should not be expected to reduce child exposure to fecal contamination in other similar settings.

## Introduction

Water, sanitation, and handwashing (WSH) interventions are hypothesized to interrupt transmission of fecal pathogens in the household environment, thereby reducing the risk of diarrhea.^1^ Yet few studies have rigorously assessed the effect of WSH interventions on domestic environmental fecal contamination beyond measuring drinking water quality. Previous studies have largely used observational designs to study associations between fecal indicator bacteria in the household environment (e.g. in water, on hands, in soil, on surfaces) and the quality of water, sanitation, and handwashing infrastructure.^2–6^ However, observational studies are not ideal for inferring causal effects.

A large number of randomized controlled trials (RCTs) have assessed the effect of water treatment interventions on fecal indicator bacteria levels in drinking water,^7,8^ but few sanitation and handwashing RCTs have measured environmental indicators of fecal contamination. We identified two RCTs evaluating the effect of handwashing interventions on hand contamination, neither of which reduced bacteria levels on hands.^9,10^ A recent systematic review concluded there was no evidence that improved sanitation reduces fecal contamination levels in water, on hands, on sentinel toys, on household surfaces, or in soil, and recommended that future evaluations of sanitation interventions include environmental fecal assessments.^11^ Many of these previous studies, however, assessed interventions that were poorly delivered or poorly adopted. ^12,13^

The WASH Benefits Kenya trial was a cluster-randomized controlled trial designed to test the effects of water, sanitation, handwashing, and nutritional interventions, alone and in combination, on child diarrhea prevalence, linear growth, parasite infections, biomarkers of environmental enteric dysfunction (EED), and child development.^14^ Unlike most previous studies, this was an efficacy trial that first designed and piloted the interventions and then took special efforts to ensure coverage and use among the study population. We recently reported the trial’s primary outcomes: WSH interventions, whether separately or in combination, did not reduce child diarrhea or improve child growth during the trial.^15^ The combined WSH and single water treatment interventions reduced roundworm (*Ascaris lumbricoides*) infection prevalence, but did not affect *Giardia* infection prevalence.^16^

To assess the extent to which the interventions may have reduced child exposure to fecal contamination, a key intermediate step to health outcomes, we nested environmental sample collection within a subset of enrolled households in selected trial arms. Our aim was to determine if the interventions reduced levels of fecal indicator bacteria in the domestic environment along likely exposure pathways for young children.

## Methods

The main trial protocol and study design has been published elsewhere.^14^ The trial enrolled pregnant women in the Kakamega, Bungoma, and Vihiga counties of Western Kenya and followed their newborns (index children) for their first two years of life. The intervention arms included water treatment, sanitation, handwashing with soap, combined WSH, nutrition, and combined WSH and nutrition (WSHN). The WSH interventions were designed to improve the environmental conditions of the children in the first two years of life to reduce early-life exposure to fecal pathogens. The water treatment intervention included installation of chlorine dispensers at community water locations plus bottled chlorine provided to enrolled households. The sanitation intervention included pit latrine upgrades with a reinforced slab and drop hole cover, as well as child potties and scoops to remove human or animal feces from households and compounds. The handwashing intervention included dual dispenser tippy-tap devices operated with independent pedals, each providing one container for soapy water^17^ (soap delivered by community promoters) and one container for rinse water, installed near the latrine and kitchen area. The nutrition intervention included lipid-based nutrient supplements (LNS) and age-appropriate recommendations on maternal nutrition and infant feeding practices. Local promoters visited study compounds at least every other month to deliver relevant behavior change messages for the treatment group assigned.

Approximately one year and two years after intervention delivery (timed to match the collection of the trial’s primary child health outcomes), we assessed environmental contamination in a subset of approximately 1,500 households that were enrolled in the EED cohort in the trial (approximately 375 children from each of the control, nutrition, combined WSH, and combined WSHN arms that participated in EED biomarker measurement). We also measured selected indicators of environmental contamination in similar sized subsets of households enrolled in the single sanitation, water, and handwashing arms. Considering improved nutrition would not be expected to influence environmental contamination, we grouped measurements collected from the WSH and WSHN arms together (WSH/WSHN) as well as measurements collected from the nutrition and control arms together (C/N). Stored drinking water samples were collected from the C/N, WSH/WSHN, water, and handwashing arms; hand rinse and toy rinse samples were collected from C/N and WSH/WSHN arms; fly densities were measured in the C/N, WSH/WSHN, and sanitation arms; mother and child visible hand cleanliness were measured in the C/N and WSH/WSHN arms (Figure 1). The intensive environmental contamination assessment was conducted in the EED subgroup because of the added benefit of being able to assess relationships between environmental contamination and EED biomarkers, as well as leverage the data collection infrastructure in the cohort. Child hand rinse and toy ball rinses (after 24 hours of play) were collected as proxy indicators of overall environmental fecal contamination in the household, and because they represent likely exposure pathways for children under two years old.^18^ Stored drinking water was sampled in the H arm because previous evidence suggested that caregiver hand and stored water contamination are highly correlated and stored drinking water fecal bacteria levels are typically less variable than hand rinse fecal bacteria levels.^4,19,20^

**Figure 1.**
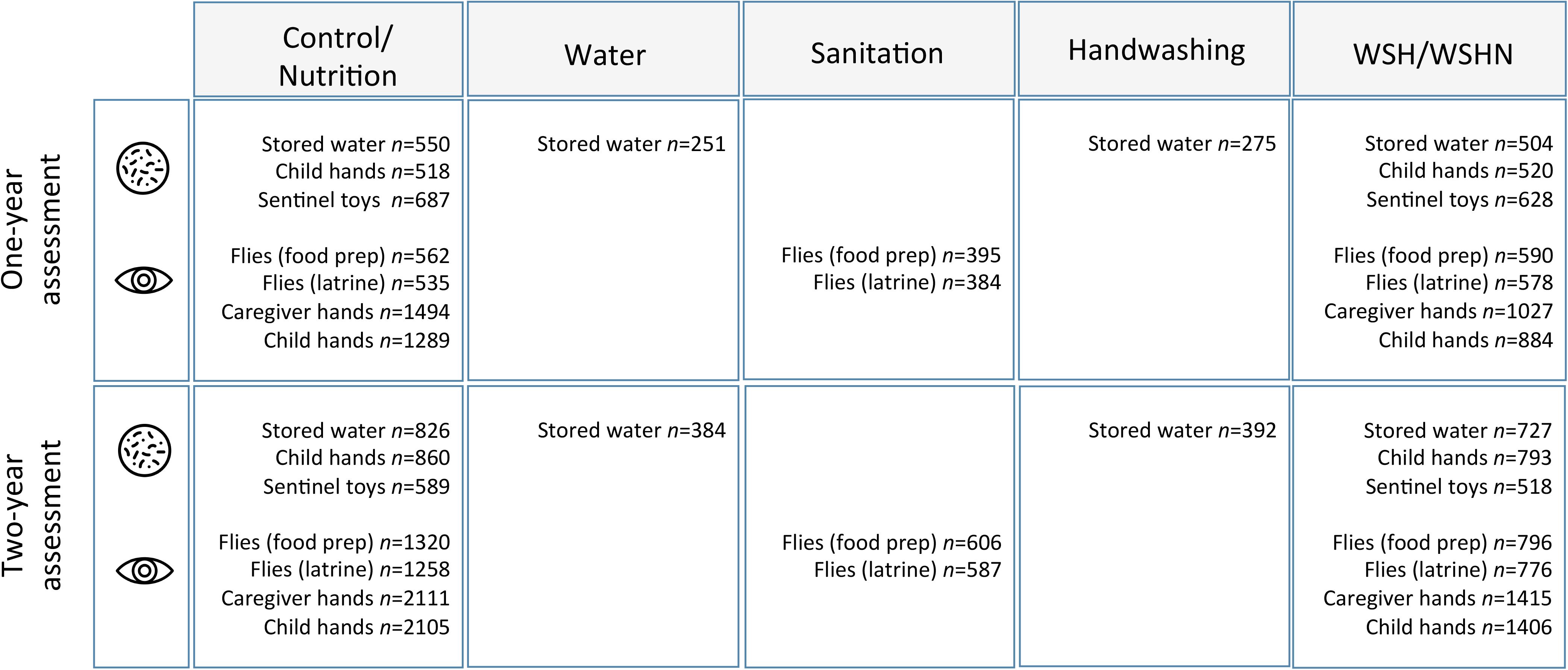
Environmental sampling profile in the trial, by study arm and year of measurement. Bacteria plate (circle with dots) indicates *E.* coli measurements; eye indicates rapid observations by field staff. The majority of measurements were conducted in the control and nutrition (C/N) and combined water, sanitation, and handwashing study arms (WSH/WSHN).

Household stored drinking water, index child hand rinse, and sentinel toy object (plastic ball) rinse samples were collected following previously published protocols^4,21^ and analyzed by membrane filtration with MI media to detect *E. coli*. Fly densities (counts) were measured at the food preparation area and at latrines using the scudder fly grill method; flies were classified as house, bottle, flesh or “other” species.^22^ Field staff observed visible cleanliness of mother and child hands (palms, fingerpads, and underneath fingernails).^4,23^ See supplemental information (SI) for additional methods on all types of sample collection and processing.

### Statistical analyses

We published a pre-specified statistical analysis plan prior to starting data analysis (https://osf.io/eg2rc/). All statistical analyses were independently replicated by two different authors (AJP, JS). Outcomes measured in the single intervention arms (water, sanitation, handwashing) and pooled combined arms (WSH/WSHN) were compared to outcomes in the pooled together C/N arms. We estimated unadjusted and adjusted intention-to-treat effects between arms, relying on the unadjusted analysis as our primary analysis. We estimated log reductions, prevalence ratios, and prevalence differences using generalized linear models with robust standard errors. We used modified Poisson regression for binary outcomes.^24,25^ All models included fixed effects for randomization block (to take advantage of the pair-matched design). In adjusted analyses, we included pre-specified variables strongly associated with the outcome to potentially improve the precision of our estimates (see SI for further details and adjusted results). We conducted subgroup analyses by year of data collection (year 1 versus year 2).

## Results and Discussion

We previously reported indicators of intervention uptake.^15^ In water intervention arms, the proportion of households with detectable chlorine residual in their stored drinking water ranged from 39 to 43% at year one and from 19 to 23% at year two. Among households in the sanitation intervention arms, the proportion with access to an improved latrine was 89-90% at year one and 78-82% at year two. In handwashing intervention arms, soap and water was present at a handwashing location at 76-78% of households at year one and at 19-23% at year two. We collected and measured levels of fecal indicator bacteria in 3909 drinking water samples, 2691 child hand rinses, and 2422 child toy ball rinses (including both the one- and two-year assessments). We collected 9653 caregiver and 9020 child hand cleanliness observations. Flies were counted at 4269 food preparation areas and 4118 latrines. See Figure 1 for samples sizes by treatment status.

### Mean levels of environmental contamination in the control group

When combining the one- and two-year assessments, 94% of stored drinking water samples were contaminated with *E. coli* (log_10_ mean 1.48 colony forming units [CFU]/100 ml), 90% of child hands were contaminated with *E. coli* (log_10_ mean 1.74 CFU/two-hands), and 73% of toys were contaminated with *E. coli* (log_10_ mean 0.58 CFU/toy) in the control group (C/N). One quarter (26%) of caregivers had visible dirt observed on their palms or fingerpads, and over half (54%) had dirt observed underneath their fingernails. Approximately one third (36%) of children had visible dirt observed on their palms or fingerpads, while two thirds (67%) had dirt observed underneath their fingernails. Flies were present at 62% of food preparation areas (mean 3.4 flies, SD 6.2) and at 67% of latrines (mean 3.7 flies, SD 5.5). House flies were more common at food preparation areas (60% prevalence) than latrines (30%); bottle flies were observed at 3% of food preparation areas but were observed at more than half of latrines (55% prevalence); flesh flies were observed at <1% of food preparation areas and 3% of latrines.

### Intervention effects on water quality

Water treatment reduced the presence of *E. coli* in drinking water by 34% (Prevalence Ratio[PR]: 0.66, 95% Confidence interval[CI] 0.58, 0.76) after one year of intervention exposure (Figure 2) and by 24% (PR: 0.76, 95% CI 0.72, 0.81) two years after interventions began (Figure 3). The combined WSH intervention showed similar effects on water quality as water treatment alone (45% reduction in *E. coli* at year 1; 19% reduction at year 2). Our findings confirm that chlorination is an effective method to improve drinking water microbial quality in low-income settings.^8^ If adoption of chlorine had been higher in the trial, the reductions in *E. coli* contamination likely would have been larger. ^26,27^ The effectiveness of the water intervention in improving water quality is consistent with observed reductions in roundworm (*Ascaris lumbricoides)* infection prevalence among children receiving the water intervention in the trial.^28^ *Ascaris* infection prevalence was reduced by 18-22% in the intervention arms that included a water treatment component (water, WSH, and WSHN arms), but not in other intervention arms.

**Figure 2.**
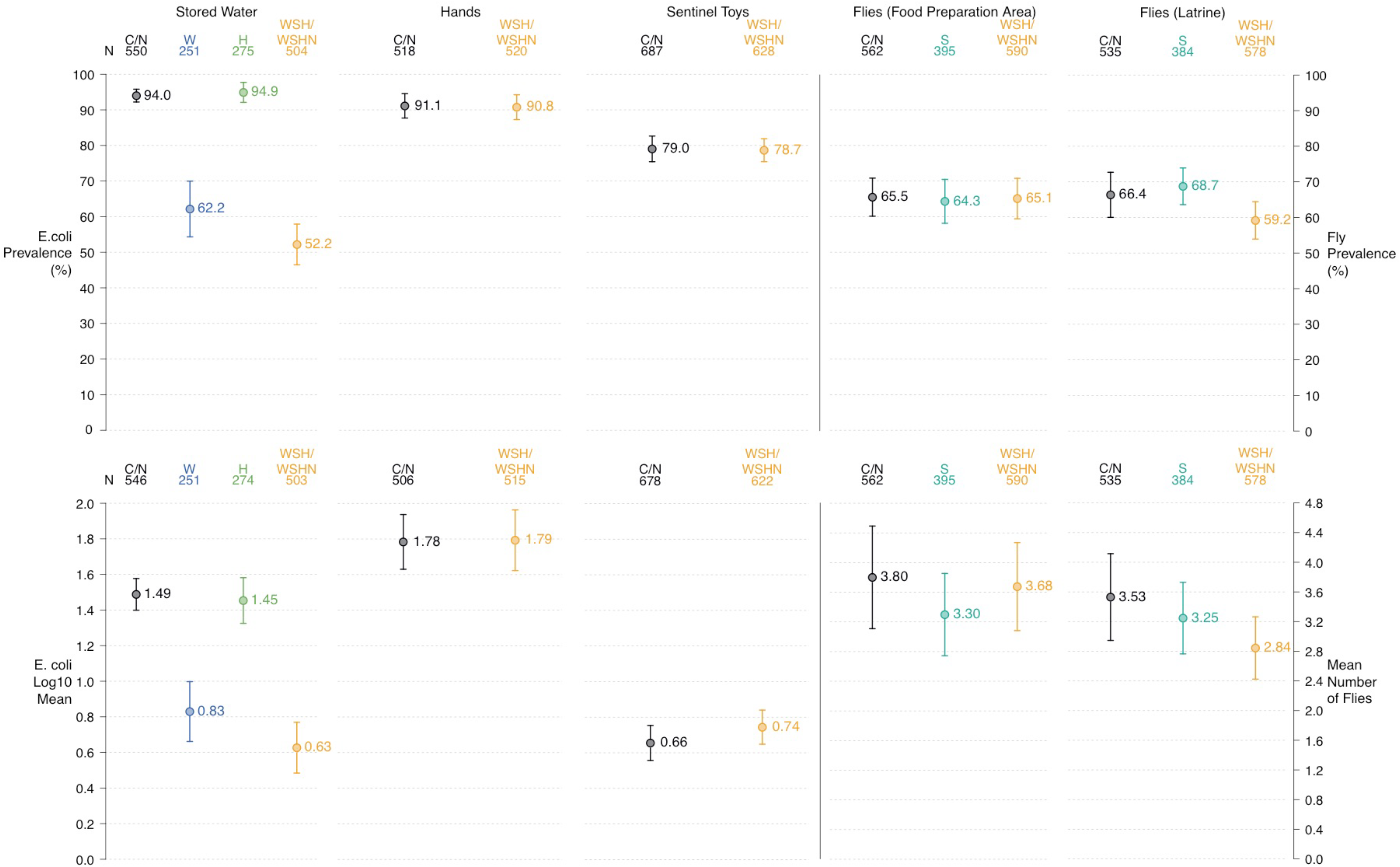
One-year assessment. Prevalence and mean concentrations of *E. coli* in stored water, on child hands, and on child toys by study arm (left); prevalence and concentration of flies measured at the food preparation and latrine areas by study arm(right).

**Figure 3.**
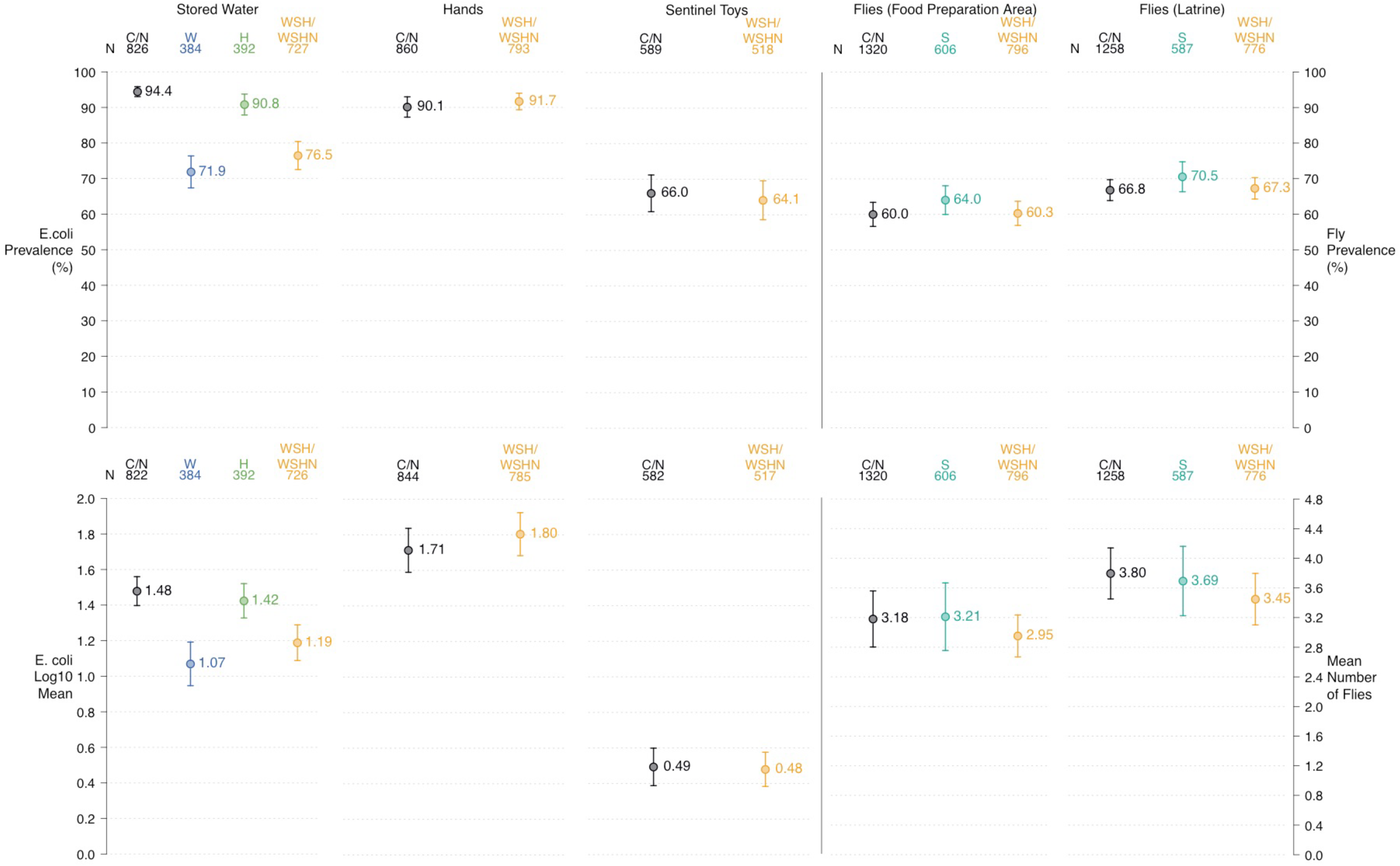
Two-year assessment. Prevalence (%) and mean concentrations of *E. coli* in stored water (log mean CFU/100ml), on child hands (log mean CFU/two hands), and on toy balls (log mean CFU/toy) by study arm (left); prevalence and concentration of flies measured at the food preparation and latrine areas by study arm (right).

Handwashing with soap slightly reduced the prevalence of *E. coli* contamination in stored drinking water at year two (PR: 0.95, 95% CI 0.92, 0.99), but not at year one. A previous study in Tanzania found that the level of fecal indicator bacteria on caregiver hands was the strongest predictor of the level of fecal indicator bacteria in stored drinking water.^4^ Our results support a link between hand contamination and stored drinking water quality; however, the magnitude of the improvement in stored drinking water quality was small. Further trials that achieve high handwashing with soap rates could be useful to better quantify the effect of increased frequency of handwashing with soap on stored drinking water quality.

### Interventions effects on child hand and toy contamination

The WSH intervention did not affect the presence or levels of fecal indicator bacteria on child hands or toy balls at any time point (Figures 2 and 3). Our findings are consistent with the parallel WASH Benefits trial in Bangladesh, which also reported no reduction in child hand or toy *E. coli* contamination in the WSH intervention arms (Ercumen et al., in press). Although it’s possible that substantial animal fecal contamination masked reductions in human fecal contamination along these pathways, a previous study found no effect of the WASH Benefits Bangladesh sanitation intervention on human host-specific fecal markers.^29^ Even if the majority of *E. coli* detected was from non-human sources, ruminant and avian feces can contain pathogenic *E. coli* strains and other common human pathogens such as *Campylobacter*, *Giardia, and Cryptosporidium*.^30,31^

The WSH intervention reduced the prevalence of visible dirt on caregiver hands (PR averaged over both measurements: 0.86, 95% CI 0.78, 0.95) and under caregiver fingernails (PR over both measurements: 0.90, 95% CI 0.85, 0.96), but did not affect visible dirt on child hands or underneath child fingernails (Table S3). These data suggest handwashing frequency marginally increased in the WSH arms, but these increases were not sufficient to reduce *E. coli* contamination on hands.

An important limitation of this work is that fecal indicator bacteria can be a poor proxy for enteric pathogens. Future evaluations should consider measuring specific human pathogens in water and in the environment to better understand how WSH interventions affect transmission pathways.

### Intervention effects on fly presence and densities

The WSH intervention reduced the prevalence and density of flies near the latrine at year one (PR: 0.88, 95% CI 0.79, 0.97; −0.6 less flies counted, 95% CI −1.2, 0.0), but not at year two (Figures 2 and 3, Table S3). No interventions affected the prevalence or density of flies at the food preparation area (Figures 2 and 3). We also did not detect any differences in prevalence of fly species between the study arms (data not shown). A trial evaluating community-led total sanitation in Mali detected a reduction in fly presence at latrines,^32^ while other trials in rural India and The Gambia found no reduction.^12,33^ Fly reductions in this trial may have been limited because the intervention was delivered at the compound level and not the community level (neighboring compounds to the study compound without a pregnant woman did not receive upgraded pit latrines with covers).

### Implications for policy and practice

Promotion of chlorine improved drinking water quality, but adoption of the water treatment intervention was much lower than expected by the end of the study, emphasizing the difficulty of achieving sustained and consistent usage of household water treatment products with monthly or less frequent behavior promotion visits.^34,35^ Importantly, the combined WSH intervention did not further improve water quality over water treatment alone. We found no evidence that combining water, sanitation, and hygiene interventions led to larger reductions in fecal contamination in the household environment than single interventions, a finding consistent with no additive benefit on health outcomes measured in this trial or in other studies.^15,36–40^

Our results indicate that the intensive WSH interventions implemented in this study did not reduce levels of fecal indicator bacteria on child hands or toys, while they slightly reduced fly presence near latrines and marginally improved visible hand cleanliness of caregivers. The lack of effect on fecal indicator bacteria measured on hands and toys has several possible explanations, including inconsistent compliance of the targeted hygiene and sanitation behaviors, failure of these specific types of WSH interventions to reduce human fecal contamination on child hands and toys, or animal fecal contamination in the household environment.^29,41,42^ The failure of the interventions to reduce fecal contamination along important exposure pathways in the household is consistent with the suggests that WSH programs that aim to improve child health may need to consider interventions that cost more but also more comprehensively reduce fecal contamination in the household setting; our findings provide additional support for transformative WSH interventions.^43^

