## Supplemental Information for "Can individual and integrated water, sanitation, and handwashing interventions reduce fecal contamination in the household environment? Evidence from the WASH Benefits cluster-randomized trial in rural Kenya"

### Methods

Study Design

Clusters were defined as one or two, and in rare cases three, adjoining villages with at least six eligible pregnant women. Nine clusters were used to form a geographic block; clusters were randomized into the nine study arms within each block (geographically pair-matched randomization increases the efficiency of effect estimates as study outcomes are expected to be correlated with geographical characteristics). The EED subset of the trial for which the one- and two-year environmental assessments were conducted consists of 54 clusters per arm in four study arms, with seven households per cluster. This results in a total sample size of 1500 households for the environmental assessment (375 households per arm), where 750 households (WSH and nutrition plus WSH arms) receive the combined WSH intervention and 750 households (control and nutrition arms) serve as comparison. We assumed an intra-class correlation coefficient of 0.2 based on a previous large-scale evaluation in rural Bangladesh and a design effect of 2.2.[44] With this sample size of 750 households per comparison group, we expected the following minimum detectable effect sizes in intervention (WSH and nutrition plus WSH pooled) vs. comparison (control and nutrition pooled) groups: (1) 0.2 $\log_{10}$ reduction in *E. coli* concentration in stored water, hand rinse and toy rinse samples and (2) 7.6 flies difference from 11.5 flies in control in the latrine and food preparation areas.

We collected samples of stored drinking water, index child hand rinses, and sentinel toy object rinses, as well as enumerated and speciated flies caught in the food preparation and latrine areas of the compound. We also recorded the presence of visible dirt on children's and caregivers' hands, including fingernails, fingerpads and palms. We collected stored drinking water samples from a subset of households in the water treatment single arm (those geographically matched with EED enrolled clusters) and from a subset of households in the handwashing single arm (those geographically matched with EED enrolled clusters). These subsets were selected based on household enrollment in the same randomization block as the EED cohort. We calculated fly densities at the latrine and at the food preparation areas in a subset of households in the sanitation single arm enrolled into the same blocks as the EED cohort (Figure 1).

Sample Collection

To sample drinking water, field workers asked the respondent to provide a glass of water as if giving it to their <5 children, and recorded whether it had been obtained directly from the drinking water source or from a storage container. Field staff asked the respondent to pour the water from the glass into a sterile Whirlpak bag. If the respondent reported treating the water with chlorine, a Whirlpak bag with sodium thiosulfate tablets was used to neutralize the residual chlorine. Stored water field blanks were collected by pouring a bottle of distilled water (prepared in the lab and carried to the field in a cooler) into a Whirlpak collection bag in the field. To collect index child hand rinse samples, field workers asked the respondent to place the index child's left hand into a sterile Whirlpak bag pre-filled with 250 mL of distilled water. The hand was massaged from the outside of the bag for 15 seconds, followed by 15 seconds of shaking. The procedure was repeated with the right hand in the same bag, and the rinse water was preserved in the Whirlpak bag. Hand rinse field blanks were collected by opening and closing an unused hand rinse Whirlpak bag in the field. Field workers provided a previously sterilized plastic toy ball to the index child and encouraged the respondent to prompt the child to play with the ball. Field workers returned to the households 24 hours later to collect a rinse sample from the ball by placing the ball into a sterile Whirlpak bag pre-filled with 300 mL of distilled water and massaging it from the outside of the bag for 15 seconds, followed by 15 seconds of shaking. Toy rinse field blanks were collected by opening and closing an unused toy rinse Whirlpak bag in the field. Field workers identified a suitable location in the food preparation area (away from smoke) and latrine area (under a roof or protected from rain). At each location, they placed one scudder fly grill on the ground and took two 90 second measurements within 30 seconds of each other to record the number of flies landing on the grill, as well as categorize the captured flies as house flies (*Musca domestica*, *Fannia canicularis*, *stomoxys*), bottle flies (*Lucilia*, *chrysomya*, *calliphora*), flesh flies (*Sarcophaga*), or other flies using a visual identification chart.

Sample analysis

All water, hand rinse, and toy rinse samples were preserved on ice in coolers and transported to the field laboratory to be processed on the same day, typically within 8 hours of collection. 100ml was filtered per stored water sample and stored water field blank sample, 50ml filtered per hand rinse sample and hand rinse field blank sample, and 100ml filtered per toy rinse sample and toy rinse field blank sample. Lab technicians targeted processing 5% duplicates for each sample type by filtering the same volume from the same collection bag. Lab blanks were also processed daily by each lab technician by filtering 50mL of distilled water. Stored water and hand rinse samples were incubated at 35 °C, while toy samples were incubated at 44.5 °C. Previous work demonstrated fecal coliforms incubated at 44.5°C on toy balls to be an indicator of environmental contamination associated with sanitation conditions.[21] *E. coli* was the primary indicator of microbial contamination (reported in main text); total and fecal coliform counts were secondary indicators (reported in SI only).

Statistical analyses

Post-incubation, samples classified as “too numerous to count” were assigned a value of 500 colonies per plate. Colony forming units (CFU) of *E. coli* were $\log_{10}$ transformed for analysis of continuous variables. Samples below the detection limit were substituted with half the lower detection limit (0.5 colonies per plate) prior to log transformation.

Our primary outcomes included *E. coli* prevalence and concentration in water, hand rinse, and toy samples, and the presence and number of flies at food preparation and latrine areas. Secondary outcomes included fecal coliform prevalence and concentration in water, hand rinse, and toy samples and the percentage of children and caregivers with visible dirt on hands. We report subgroup analyses by year of data collection (year 1 versus year 2) in the main test and by season (wet vs. dry season) in the SI only.

We pre-screened covariates to assess whether they are associated with each outcome prior to including them in adjusted statistical models (Pocock et al. 2002). We used the likelihood ratio test to assess the

association between each outcome and each covariate and included covariates with a $p$-value < 0.2 in the adjusted analysis. We also excluded covariates that have little variation in the study population (e.g., prevalence <5 %).
The following baseline covariates were tested for inclusion in the adjusted models:

- ID of the lab staff member who performed the lab analysis
- Month of measurement, to account for seasonal variation
- Most recent time it rained (now, today, yesterday, day before yesterday, last week, or before last week as reported by respondent)
- Child age (days)
- Child sex
- Mother's age (years)
- Mother's education level (no education, completed primary, completed secondary)
- Number of children <18 years in the household
- Number of individuals living in the compound
- Distance (in minutes) to the household's primary drinking water source
- Housing materials (floor, walls, roof) and household assets
    - Assets measured: electricity, radio, television, mobile phone, clock, bicycle, motorcycle, stove, gas cooker, car. Number of: cows, goats, dogs, chickens.

Additionally, we tested the following covariates that would be expected to affect fecal indicator bacteria levels for these specific types of samples. While these covariates were measured at the time of the environmental assessments (rather than at baseline), we did not expect them to be impacted by the study interventions.

*Sentinel toy objects*

- Location of toy ball at time of sample collection (inside vs. outside)
- How much children played with ball (4-level categories [several times (4+), few times (2-3), once, never]
- Whether children from other compounds played with ball
- Where children played with ball (indoors vs. outdoors)
- Whether respondent cleaned ball prior to sample collection
- Condition of ball at time of sample collection (intact/bulging vs. deflated/burst/torn)

*Flies*

- Location of food preparation area (indoors vs. outdoors)

*Results – blanks and duplicates*
[to be added]

**Table S1.** *E. coli* measured at one- and two-year assessments, combined and separately, interventions vs. control.

**One- and Two-Year Assessments Combined**

| | | | | Prevalence Ratio | | | | | | Log10 Difference | | | | | |
|---|---|---|---|---|---|---|---|---|---|---|---|---|---|---|---|
| | | | | Unadjusted | | | Adjusted | | | Unadjusted | | | Adjusted | | |
| Arms | N | Prev. | Log10 Mean (CFU/ 100 mL) | Prev. Ratio | 95% Confidence Intervals | | Prev. Ratio | 95% Confidence Intervals | | Log10 Diff. | 95% Confidence Intervals | | Log10 Diff. | 95% Confidence Intervals | |
| **Stored Water** | | | | | | | | | | | | | | | |
| Control/Nutrition | 1376 | 0.94 | 1.48 | - | - | - | - | - | - | - | - | - | - | - | - |
| WSH/WSH+Nutrition | 1231 | 0.67 | 0.96 | 0.70 | 0.67 | 0.74 | 0.70 | 0.66 | 0.74 | -0.53 | -0.63 | -0.43 | -0.53 | -0.63 | -0.43 |
| Water | 635 | 0.68 | 0.97 | 0.73 | 0.68 | 0.78 | 0.72 | 0.68 | 0.78 | -0.51 | -0.62 | -0.40 | -0.49 | -0.61 | -0.38 |
| Handwashing | 667 | 0.93 | 1.44 | 0.98 | 0.95 | 1.00 | 0.98 | 0.95 | 1.00 | -0.06 | -0.16 | 0.03 | -0.10 | -0.20 | -0.01 |
| **Child Hands** | | | | | | | | | | | | | | | |
| Control/Nutrition | 1378 | 0.90 | 1.74 | - | - | - | - | - | - | - | - | - | - | - | - |
| WSH/WSH+Nutrition | 1313 | 0.91 | 1.80 | 1.00 | 0.98 | 1.03 | 1.00 | 0.98 | 1.03 | 0.04 | -0.04 | 0.13 | 0.08 | -0.01 | 0.17 |
| **Sentinel Toys** | | | | | | | | | | | | | | | |
| Control/Nutrition | 1276 | 0.73 | 0.58 | - | - | - | - | - | - | - | - | - | - | - | - |
| WSH/WSH+Nutrition | 1146 | 0.72 | 0.62 | 0.97 | 0.93 | 1.02 | 0.97 | 0.91 | 1.03 | 0.02 | -0.07 | 0.10 | -0.02 | -0.11 | 0.07 |

**One-Year Assessment**

| | | | | Prevalence Ratio | | | | | | Log10 Difference | | | | | |
|---|---|---|---|---|---|---|---|---|---|---|---|---|---|---|---|
| | | | | Unadjusted | | | Adjusted | | | Unadjusted | | | Adjusted | | |
| Arms | N | Prev. | Log10 Mean (CFU/ 100 mL) | Prev. Ratio | 95% Confidence Intervals | | Prev. Ratio | 95% Confidence Intervals | | Log10 Diff. | 95% Confidence Intervals | | Log10 Diff. | 95% Confidence Intervals | |
| **Stored Water** | | | | | | | | | | | | | | | |
| Control/Nutrition | 550 | 0.94 | 1.49 | - | - | - | - | - | - | - | - | - | - | - | - |
| WSH/WSH+Nutrition | 504 | 0.52 | 0.63 | 0.55 | 0.48 | 0.61 | 0.54 | 0.48 | 0.61 | -0.90 | -1.07 | -0.74 | -0.87 | -1.03 | -0.70 |
| Water | 251 | 0.62 | 0.83 | 0.66 | 0.58 | 0.76 | 0.65 | 0.57 | 0.75 | -0.64 | -0.86 | -0.43 | -0.66 | -0.89 | -0.42 |
| Handwashing | 275 | 0.95 | 1.45 | 1.01 | 0.97 | 1.05 | 1.02 | 0.98 | 1.05 | -0.05 | -0.22 | 0.12 | -0.08 | -0.26 | 0.09 |
| **Child Hands** | | | | | | | | | | | | | | | |
| Control/Nutrition | 518 | 0.91 | 1.78 | - | - | - | - | - | - | - | - | - | - | - | - |
| WSH/WSH+Nutrition | 520 | 0.91 | 1.79 | 1.00 | 0.95 | 1.04 | 1.00 | 0.95 | 1.04 | 0.00 | -0.16 | 0.17 | 0.07 | -0.10 | 0.23 |
| **Sentinel Toys** | | | | | | | | | | | | | | | |
| Control/Nutrition | 687 | 0.79 | 0.66 | - | - | - | - | - | - | - | - | - | - | - | - |
| WSH/WSH+Nutrition | 628 | 0.79 | 0.74 | 0.98 | 0.93 | 1.04 | 0.98 | 0.89 | 1.07 | 0.06 | -0.08 | 0.19 | 0.02 | -0.13 | 0.16 |

**Two-Year Assessment**

| | | | | Prevalence Ratio | | | | | | Log10 Difference | | | | | |
|---|---|---|---|---|---|---|---|---|---|---|---|---|---|---|---|
| | | | | Unadjusted | | | Adjusted | | | Unadjusted | | | Adjusted | | |
| Arms | N | Prev. | Log10 Mean (CFU/ 100 mL) | Prev. Ratio | 95% Confidence Intervals | | Prev. Ratio | 95% Confidence Intervals | | Log10 Diff. | 95% Confidence Intervals | | Log10 Diff. | 95% Confidence Intervals | |
| **Stored Water** | | | | | | | | | | | | | | | |
| Control/Nutrition | 826 | 0.94 | 1.48 | - | - | - | - | - | - | - | - | - | - | - | - |
| WSH/WSH+Nutrition | 727 | 0.76 | 1.19 | 0.81 | 0.76 | 0.85 | 0.81 | 0.76 | 0.85 | -0.29 | -0.40 | -0.18 | -0.30 | -0.41 | -0.19 |
| Water | 384 | 0.72 | 1.07 | 0.76 | 0.72 | 0.81 | 0.77 | 0.72 | 0.82 | -0.43 | -0.54 | -0.32 | -0.37 | -0.49 | -0.25 |
| Handwashing | 392 | 0.91 | 1.42 | 0.95 | 0.92 | 0.99 | 0.95 | 0.92 | 0.98 | -0.07 | -0.19 | 0.05 | -0.10 | -0.22 | 0.03 |
| **Child Hands** | | | | | | | | | | | | | | | |
| Control/Nutrition | 860 | 0.90 | 1.71 | - | - | - | - | - | - | - | - | - | - | - | - |
| WSH/WSH+Nutrition | 793 | 0.92 | 1.80 | 1.01 | 0.98 | 1.04 | 1.01 | 0.98 | 1.04 | 0.07 | -0.05 | 0.20 | 0.13 | 0.00 | 0.26 |
| **Sentinel Toys** | | | | | | | | | | | | | | | |
| Control/Nutrition | 589 | 0.66 | 0.49 | - | - | - | - | - | - | - | - | - | - | - | - |
| WSH/WSH+Nutrition | 518 | 0.64 | 0.48 | 0.96 | 0.88 | 1.04 | 0.97 | 0.89 | 1.07 | -0.04 | -0.15 | 0.07 | -0.04 | -0.15 | 0.06 |

**Table S2.** Total coliforms (water and hand samples) or fecal coliforms (toy samples) measured at one- and two-year assessments, combined and separately, interventions vs. control.

**One- and Two-Year Assessments Combined**

| | | | | Prevalence Ratio | | | | | | Log10 Difference | | | | | |
|---|---|---|---|---|---|---|---|---|---|---|---|---|---|---|---|
| | | | | Unadjusted | | | Adjusted | | | Unadjusted | | | Adjusted | | |
| Arms | N | Prev. | Log10 Mean (CFU/ 100 mL) | Prev. Ratio | 95% Confidence Intervals | | Prev. Ratio | 95% Confidence Intervals | | Log10 Diff. | 95% Confidence Intervals | | Log10 Diff. | 95% Confidence Intervals | |
| **Stored Water** | | | | | | | | | | | | | | | |
| Control/Nutrition | 1376 | 0.99 | 2.56 | - | - | - | - | - | - | - | - | - | - | - | - |
| WSH/WSH+Nutrition | 1231 | 0.81 | 1.80 | 0.82 | 0.79 | 0.85 | 0.82 | 0.79 | 0.85 | -0.76 | -0.85 | -0.67 | -0.75 | -0.83 | -0.66 |
| Water | 635 | 0.84 | 1.85 | 0.85 | 0.82 | 0.89 | 0.85 | 0.82 | 0.89 | -0.69 | -0.83 | -0.56 | -0.66 | -0.79 | -0.54 |
| Handwashing | 664 | 0.98 | 2.52 | 0.99 | 0.98 | 1.00 | 0.99 | 0.98 | 1.01 | -0.03 | -0.09 | 0.03 | -0.03 | -0.09 | 0.02 |
| **Child Hands** | | | | | | | | | | | | | | | |
| Control/Nutrition | 1376 | 0.99 | 2.71 | - | - | - | - | - | - | - | - | - | - | - | - |
| WSH/WSH+Nutrition | 1312 | 0.99 | 2.72 | 1.00 | 0.99 | 1.01 | 1.00 | 0.99 | 1.01 | -0.01 | -0.05 | 0.04 | -0.01 | -0.06 | 0.04 |
| **Sentinel Toys** | | | | | | | | | | | | | | | |
| Control/Nutrition | 1276 | 0.90 | 1.23 | - | - | - | - | - | - | - | - | - | - | - | - |
| WSH/WSH+Nutrition | 1146 | 0.90 | 1.25 | 0.98 | 0.96 | 1.01 | 0.98 | 0.94 | 1.02 | -0.01 | -0.09 | 0.07 | -0.02 | -0.11 | 0.07 |

**One-Year Assessment**

| | | | | Prevalence Ratio | | | | | | Log10 Difference | | | | | |
|---|---|---|---|---|---|---|---|---|---|---|---|---|---|---|---|
| | | | | Unadjusted | | | Adjusted | | | Unadjusted | | | Adjusted | | |
| Arms | N | Prev. | Log10 Mean (CFU/ 100 mL) | Prev. Ratio | 95% Confidence Intervals | | Prev. Ratio | 95% Confidence Intervals | | Log10 Diff. | 95% Confidence Intervals | | Log10 Diff. | 95% Confidence Intervals | |
| **Stored Water** | | | | | | | | | | | | | | | |
| Control/Nutrition | 550 | 0.99 | 2.54 | - | - | - | - | - | - | - | - | - | - | - | - |
| WSH/WSH+Nutrition | 504 | 0.71 | 1.34 | 0.71 | 0.67 | 0.77 | 0.71 | 0.67 | 0.77 | -1.22 | -1.36 | -1.08 | -1.17 | -1.31 | -1.03 |
| Water | 251 | 0.82 | 1.66 | 0.82 | 0.77 | 0.87 | 0.82 | 0.76 | 0.87 | -0.87 | -1.09 | -0.65 | -0.86 | -1.10 | -0.63 |
| Handwashing | 272 | 0.98 | 2.50 | 0.99 | 0.97 | 1.01 | 0.99 | 0.97 | 1.01 | -0.03 | -0.10 | 0.04 | -0.01 | -0.09 | 0.07 |
| **Child Hands** | | | | | | | | | | | | | | | |
| Control/Nutrition | 516 | 0.99 | 2.63 | - | - | - | - | - | - | - | - | - | - | - | - |
| WSH/WSH+Nutrition | 520 | 0.99 | 2.62 | 1.00 | 0.98 | 1.01 | 1.00 | 0.98 | 1.01 | -0.02 | -0.10 | 0.06 | -0.01 | -0.08 | 0.07 |
| **Sentinel Toys** | | | | | | | | | | | | | | | |
| Control/Nutrition | 687 | 0.93 | 1.36 | - | - | - | - | - | - | - | - | - | - | - | - |
| WSH/WSH+Nutrition | 628 | 0.93 | 1.38 | 0.98 | 0.94 | 1.02 | 0.97 | 0.92 | 1.03 | -0.01 | -0.14 | 0.12 | 0.03 | -0.12 | 0.18 |

**Two-Year Assessment**

| | | | | Prevalence Ratio | | | | | | Log10 Difference | | | | | |
|---|---|---|---|---|---|---|---|---|---|---|---|---|---|---|---|
| | | | | Unadjusted | | | Adjusted | | | Unadjusted | | | Adjusted | | |
| Arms | N | Prev. | Log10 Mean (CFU/ 100 mL) | Prev. Ratio | 95% Confidence Intervals | | Prev. Ratio | 95% Confidence Intervals | | Log10 Diff. | 95% Confidence Intervals | | Log10 Diff. | 95% Confidence Intervals | |
| **Stored Water** | | | | | | | | | | | | | | | |
| Control/Nutrition | 826 | 0.99 | 2.57 | - | - | - | - | - | - | - | - | - | - | - | - |
| WSH/WSH+Nutrition | 727 | 0.88 | 2.11 | 0.89 | 0.86 | 0.92 | 0.89 | 0.86 | 0.92 | -0.46 | -0.56 | -0.36 | -0.46 | -0.57 | -0.36 |
| Water | 384 | 0.86 | 1.97 | 0.87 | 0.83 | 0.92 | 0.87 | 0.83 | 0.92 | -0.59 | -0.74 | -0.43 | -0.55 | -0.71 | -0.39 |
| Handwashing | 392 | 0.99 | 2.54 | 1.00 | 0.98 | 1.01 | 1.00 | 0.98 | 1.01 | -0.04 | -0.13 | 0.04 | -0.04 | -0.12 | 0.04 |
| **Child Hands** | | | | | | | | | | | | | | | |
| Control/Nutrition | 860 | 0.99 | 2.75 | - | - | - | - | - | - | - | - | - | - | - | - |
| WSH/WSH+Nutrition | 792 | 1.00 | 2.78 | 1.00 | 1.00 | 1.01 | 1.00 | 1.00 | 1.01 | 0.00 | -0.05 | 0.05 | 0.01 | -0.05 | 0.06 |
| **Sentinel Toys** | | | | | | | | | | | | | | | |
| Control/Nutrition | 589 | 0.86 | 1.07 | - | - | - | - | - | - | - | - | - | - | - | - |
| WSH/WSH+Nutrition | 518 | 0.86 | 1.10 | 0.99 | 0.94 | 1.04 | 0.98 | 0.93 | 1.05 | -0.02 | -0.14 | 0.10 | -0.06 | -0.18 | 0.06 |

**Table S3.** Fly counts and dirt on hands/fingernails measured at one- and two-year assessments, combined and separately, interventions vs. control

**One- and Two-Year Assessments Combined**

| Arms | N | Prev. | Mean | Prevalence Ratio | | | | | | Fly Count Difference | | | | | |
|---|---|---|---|---|---|---|---|---|---|---|---|---|---|---|---|
| | | | | Unadjusted | | | Adjusted | | | Unadjusted | | | Adjusted | | |
| | | | | Prev. Ratio | 95% Confidence Intervals | | Prev. Ratio | 95% Confidence Intervals | | Fly Diff. | 95% Confidence Intervals | | Fly Diff. | 95% Confidence Intervals | |
| **Flies Caught in Food Preparation Area** | | | | | | | | | | | | | | | |
| Control/Nutrition | 1882 | 0.62 | 3.37 | - | - | - | - | - | - | - | - | - | - | - | - |
| WSH/WSH+Nutrition | 1386 | 0.62 | 3.26 | 1.01 | 0.94 | 1.08 | 1.00 | 0.94 | 1.07 | -0.12 | -0.60 | 0.36 | -0.22 | -0.70 | 0.26 |
| Sanitation | 1001 | 0.641 | 3.25 | 0.99 | 0.93 | 1.06 | 0.98 | 0.92 | 1.05 | -0.30 | -0.88 | 0.27 | -0.23 | -0.78 | 0.31 |
| **Flies Caught in Latrine Area** | | | | | | | | | | | | | | | |
| Control/Nutrition | 1793 | 0.67 | 3.72 | - | - | - | - | - | - | - | - | - | - | - | - |
| WSH/WSH+Nutrition | 1354 | 0.64 | 3.19 | 0.96 | 0.91 | 1.01 | 0.96 | 0.91 | 1.01 | -0.49 | -0.87 | -0.11 | -0.52 | -0.89 | -0.15 |
| Sanitation | 971 | 0.70 | 3.52 | 1.02 | 0.96 | 1.08 | 1.01 | 0.96 | 1.07 | -0.49 | -1.00 | 0.01 | -0.42 | -0.95 | 0.10 |
| **Visible Dirt on Caregiver Hands** | | | | | | | | | | | | | | | |
| Control/Nutrition | 3605 | 0.26 | - | - | - | - | - | - | - | - | - | - | - | - | - |
| WSH/WSH+Nutrition | 2442 | 0.23 | - | 0.86 | 0.78 | 0.95 | 0.86 | 0.75 | 0.98 | - | - | - | - | - | - |
| **Visible Dirt under Caregiver Fingernails** | | | | | | | | | | | | | | | |
| Control/Nutrition | 3605 | 0.54 | - | - | - | - | - | - | - | - | - | - | - | - | - |
| WSH/WSH+Nutrition | 2442 | 0.48 | - | 0.90 | 0.85 | 0.96 | 0.91 | 0.84 | 0.98 | - | - | - | - | - | - |
| **Visible Dirt on Child Hands** | | | | | | | | | | | | | | | |
| Control/Nutrition | 3390 | 0.36 | - | - | - | - | - | - | - | - | - | - | - | - | - |
| WSH/WSH+Nutrition | 2288 | 0.34 | - | 0.92 | 0.85 | 1.01 | 0.87 | 0.78 | 0.97 | - | - | - | - | - | - |
| **Visible Dirt under Child Fingernails** | | | | | | | | | | | | | | | |
| Control/Nutrition | 3394 | 0.67 | - | - | - | - | - | - | - | - | - | - | - | - | - |
| WSH/WSH+Nutrition | 2290 | 0.66 | - | 0.99 | 0.94 | 1.03 | 0.95 | 0.89 | 1.02 | - | - | - | - | - | - |

**One-Year Assessment**

| Arms | N | Prev. | Mean | Prevalence Ratio | | | | | | Fly Count Difference | | | | | |
|---|---|---|---|---|---|---|---|---|---|---|---|---|---|---|---|
| | | | | Unadjusted | | | Adjusted | | | Unadjusted | | | Adjusted | | |
| | | | | Prev. Ratio | 95% Confidence Intervals | | Prev. Ratio | 95% Confidence Intervals | | Fly Diff. | 95% Confidence Intervals | | Fly Diff. | 95% Confidence Intervals | |
| **Flies Caught in Food Preparation Area** | | | | | | | | | | | | | | | |
| Control/Nutrition | 562 | 0.65 | 3.80 | - | - | - | - | - | - | - | - | - | - | - | - |
| WSH/WSH+Nutrition | 590 | 0.65 | 3.68 | 0.97 | 0.89 | 1.06 | 0.98 | 0.89 | 1.07 | -0.28 | -1.18 | 0.63 | -0.24 | -1.17 | 0.69 |
| Sanitation | 395 | 0.64 | 3.30 | 0.94 | 0.81 | 1.10 | 0.93 | 0.79 | 1.09 | -0.42 | -1.50 | 0.66 | -0.69 | -1.87 | 0.49 |
| **Flies Caught in Latrine Area** | | | | | | | | | | | | | | | |
| Control/Nutrition | 535 | 0.66 | 3.53 | - | - | - | - | - | - | - | - | - | - | - | - |
| WSH/WSH+Nutrition | 578 | 0.59 | 2.84 | 0.88 | 0.79 | 0.97 | 0.86 | 0.77 | 0.95 | -0.59 | -1.16 | -0.02 | -0.66 | -1.24 | -0.08 |
| Sanitation | 384 | 0.69 | 3.25 | 0.97 | 0.84 | 1.12 | 0.94 | 0.81 | 1.10 | -0.18 | -0.92 | 0.57 | -0.55 | -1.43 | 0.33 |
| **Visible Dirt on Caregiver Hands** | | | | | | | | | | | | | | | |
| Control/Nutrition | 1494 | 0.26 | - | - | - | - | - | - | - | - | - | - | - | - | - |
| WSH/WSH+Nutrition | 1027 | 0.21 | - | 0.81 | 0.70 | 0.93 | 0.91 | 0.75 | 1.11 | - | - | - | - | - | - |
| **Visible Dirt under Caregiver Fingernails** | | | | | | | | | | | | | | | |
| Control/Nutrition | 1494 | 0.47 | - | - | - | - | - | - | - | - | - | - | - | - | - |
| WSH/WSH+Nutrition | 1027 | 0.40 | - | 0.87 | 0.79 | 0.96 | 0.91 | 0.80 | 1.05 | - | - | - | - | - | - |
| **Visible Dirt on Child Hands** | | | | | | | | | | | | | | | |
| Control/Nutrition | 1285 | 0.31 | - | - | - | - | - | - | - | - | - | - | - | - | - |
| WSH/WSH+Nutrition | 882 | 0.28 | - | 0.87 | 0.74 | 1.03 | 0.79 | 0.61 | 1.02 | - | - | - | - | - | - |
| **Visible Dirt under Child Fingernails** | | | | | | | | | | | | | | | |
| Control/Nutrition | 1289 | 0.57 | - | - | - | - | - | - | - | - | - | - | - | - | - |
| WSH/WSH+Nutrition | 884 | 0.56 | - | 1.00 | 0.91 | 1.09 | 1.01 | 0.89 | 1.15 | - | - | - | - | - | - |

| | | | | Two-Year Assessment | | | | | | | | | | | |
|---|---|---|---|---|---|---|---|---|---|---|---|---|---|---|---|
| | | | | Prevalence Ratio | | | | | | Fly Count Difference | | | | | |
| | | | | Unadjusted | | | Adjusted | | | Unadjusted | | | Adjusted | | |
| **Arms** | **N** | **Prev.** | **Mean** | **Prev. Ratio** | **95% Confidence Intervals** | | **Prev. Ratio** | **95% Confidence Intervals** | | **Fly Diff.** | **95% Confidence Intervals** | | **Fly Diff.** | **95% Confidence Intervals** | |
| **Flies Caught in Food Preparation Area** | | | | | | | | | | | | | | | |
| Control/Nutrition | 1320 | 0.60 | 3.18 | - | - | - | - | - | - | - | - | - | - | - | - |
| WSH/WSH+Nutrition | 796 | 0.60 | 2.95 | 1.01 | 0.93 | 1.10 | 1.01 | 0.93 | 1.10 | -0.20 | -0.72 | 0.33 | -0.16 | -0.67 | 0.35 |
| Sanitation | 606 | 0.64 | 3.21 | 1.01 | 0.93 | 1.09 | 1.01 | 0.93 | 1.09 | -0.24 | -0.95 | 0.47 | -0.02 | -0.64 | 0.60 |
| **Flies Caught in Latrine Area** | | | | | | | | | | | | | | | |
| Control/Nutrition | 1258 | 0.67 | 3.80 | - | - | - | - | - | - | - | - | - | - | - | - |
| WSH/WSH+Nutrition | 776 | 0.67 | 3.45 | 1.02 | 0.96 | 1.08 | 1.01 | 0.95 | 1.08 | -0.34 | -0.78 | 0.10 | -0.48 | -0.91 | -0.06 |
| Sanitation | 587 | 0.71 | 3.69 | 1.04 | 0.96 | 1.13 | 1.03 | 0.95 | 1.11 | -0.46 | -1.09 | 0.16 | -0.56 | -1.17 | 0.05 |
| **Visible Dirt on Caregiver Hands** | | | | | | | | | | | | | | | |
| Control/Nutrition | 2111 | 0.26 | - | - | - | - | - | - | - | - | - | - | - | - | - |
| WSH/WSH+Nutrition | 1415 | 0.24 | - | 0.90 | 0.79 | 1.02 | 0.84 | 0.69 | 1.01 | - | - | - | - | - | - |
| **Visible Dirt under Caregiver Fingernails** | | | | | | | | | | | | | | | |
| Control/Nutrition | 2111 | 0.58 | - | - | - | - | - | - | - | - | - | - | - | - | - |
| WSH/WSH+Nutrition | 1415 | 0.54 | - | 0.92 | 0.87 | 0.99 | 0.91 | 0.83 | 1.00 | - | - | - | - | - | - |
| **Visible Dirt on Child Hands** | | | | | | | | | | | | | | | |
| Control/Nutrition | 2105 | 0.39 | - | - | - | - | - | - | - | - | - | - | - | - | - |
| WSH/WSH+Nutrition | 1406 | 0.37 | - | 0.95 | 0.86 | 1.04 | 0.91 | 0.79 | 1.05 | - | - | - | - | - | - |
| **Visible Dirt under Child Fingernails** | | | | | | | | | | | | | | | |
| Control/Nutrition | 2105 | 0.74 | - | - | - | - | - | - | - | - | - | - | - | - | - |
| WSH/WSH+Nutrition | 1406 | 0.72 | - | 0.98 | 0.93 | 1.03 | 0.94 | 0.88 | 1.01 | - | - | - | - | - | - |

**Table S4.** *E. coli* and fly prevalence and concentration measured at one- and two-year assessments, combined and separately, interventions vs. control, subgroup analysis by season.

| One- and Two-Year Assessments Combined | | | | | | | | | | | | | | | | | | | | | |
|---|---|---|---|---|---|---|---|---|---|---|---|---|---|---|---|---|---|---|---|---|---|
| | Prevalence | | | | | | | | | | | Concentration | | | | | | | | | |
| | Wet Season | | | | | Dry Season | | | | | | | Wet Season | | | | | Dry Season | | | |
| Arms | N | Prev. | Prev. Ratio | 95% Confidence Intervals | | N | Prev. | Prev. Ratio | 95% Confidence Intervals | | Inter-action P-Value | N | Log10 Mean (CFU/ 100 mL)/ Mean (Flies) | Log Diff./ Fly Count | 95% Confidence Intervals | | N | Log10 Mean (CFU/ 100 mL)/ Mean (Flies) | Log Diff./ Fly Count | 95% Confidence Intervals | | Inter-action P-Value |
| **Stored Water** | | | | | | | | | | | | | | | | | | | | | |
| Control/Nutrition | 737 | 0.94 | - | - | - | 637 | 0.94 | - | - | - | - | 731 | 1.51 | - | - | - | 635 | 1.44 | - | - | - | - |
| WSH/ WSH+Nutrition | 665 | 0.65 | 0.69 | 0.64 | 0.75 | 565 | 0.68 | 0.72 | 0.67 | 0.77 | 0.544 | 664 | 0.95 | -0.57 | -0.70 | -0.44 | 564 | 0.966 | -0.47 | -0.61 | -0.34 | 0.282 |
| Water | 361 | 0.65 | 0.69 | 0.62 | 0.76 | 274 | 0.72 | 0.77 | 0.71 | 0.84 | 0.091 | 361 | 0.95 | -0.57 | -0.75 | -0.39 | 274 | 1.003 | -0.44 | -0.56 | -0.32 | 0.262 |
| Handwashing | 363 | 0.93 | 0.98 | 0.94 | 1.02 | 304 | 0.92 | 0.97 | 0.93 | 1.01 | 0.834 | 362 | 1.47 | -0.05 | -0.18 | 0.08 | 304 | 1.395 | -0.07 | -0.21 | 0.07 | 0.812 |
| **Child Hands** | | | | | | | | | | | | | | | | | | | | | |
| Control/Nutrition | 743 | 0.89 | - | - | - | 632 | 0.92 | - | - | - | - | 724 | 1.68 | - | - | - | 623 | 1.81 | - | - | - | - |
| WSH/ WSH+Nutrition | 714 | 0.89 | 0.99 | 0.96 | 1.03 | 597 | 0.93 | 1.02 | 0.99 | 1.04 | 0.370 | 705 | 1.71 | -0.02 | -0.13 | 0.10 | 593 | 1.905 | 0.13 | -0.01 | 0.26 | 0.127 |
| **Sentinel Toys** | | | | | | | | | | | | | | | | | | | | | |
| Control/Nutrition | 627 | 0.72 | - | - | - | 649 | 0.74 | - | - | - | - | 616 | 0.59 | - | - | - | 644 | 0.57 | - | - | - | - |
| WSH/ WSH+Nutrition | 591 | 0.73 | 0.99 | 0.93 | 1.04 | 555 | 0.71 | 0.96 | 0.89 | 1.03 | 0.534 | 587 | 0.67 | 0.02 | -0.07 | 0.11 | 552 | 0.57 | 0.01 | -0.13 | 0.15 | 0.873 |
| **Flies Caught in Food Preparation Area** | | | | | | | | | | | | | | | | | | | | | |
| Control/Nutrition | 946 | 0.65 | - | - | - | 936 | 0.59 | - | - | - | - | 946 | 3.78 | - | - | - | 936 | 2.95 | - | - | - | - |
| WSH/ WSH+Nutrition | 745 | 0.62 | 0.98 | 0.91 | 1.05 | 641 | 0.62 | 1.03 | 0.93 | 1.14 | 0.396 | 745 | 3.35 | -0.54 | -1.26 | 0.18 | 641 | 3.15 | 0.24 | -0.28 | 0.76 | 0.075 |
| Sanitation | 527 | 0.68 | 1.00 | 0.92 | 1.09 | 474 | 0.60 | 0.97 | 0.87 | 1.09 | 0.724 | 527 | 3.63 | -0.48 | -1.35 | 0.38 | 474 | 2.82 | -0.11 | -0.72 | 0.49 | 0.462 |
| **Flies Caught in Latrine Area** | | | | | | | | | | | | | | | | | | | | | |
| Control/Nutrition | 912 | 0.70 | - | - | - | 881 | 0.63 | - | - | - | - | 912 | 4.17 | - | - | - | 881 | 3.25 | - | - | - | - |
| WSH/ WSH+Nutrition | 731 | 0.64 | 0.93 | 0.87 | 1.00 | 623 | 0.63 | 0.97 | 0.89 | 1.07 | 0.488 | 731 | 3.20 | -0.82 | -1.29 | -0.34 | 623 | 3.17 | -0.22 | -0.78 | 0.34 | 0.111 |
| Sanitation | 515 | 0.70 | 0.96 | 0.89 | 1.04 | 456 | 0.69 | 1.09 | 1.00 | 1.18 | 0.041 | 515 | 3.74 | -0.84 | -1.58 | -0.09 | 456 | 3.27 | -0.12 | -0.70 | 0.46 | 0.110 |

| One-Year Assessment | | | | | | | | | | | | | | | | | | | | | |
|---|---|---|---|---|---|---|---|---|---|---|---|---|---|---|---|---|---|---|---|---|---|
| | Prevalence | | | | | | | | | | Concentration | | | | | | | | | | |
| | Wet Season | | | | | Dry Season | | | | | | Wet Season | | | | | Dry Season | | | | | |
| Arms | N | Prev. | Prev. Ratio | 95% Confidence Intervals | | N | Prev. | Prev. Ratio | 95% Confidence Intervals | | Inter-action P-Value | N | Log10 Mean (CFU/ 100 mL)/ Mean (Flies) | Log Diff./ Fly Count | 95% Confidence Intervals | | N | Log10 Mean (CFU/ 100 mL)/ Mean (Flies) | Log Diff./ Fly Count | 95% Confidence Intervals | | Inter-action P-Value |
| **Stored Water** | | | | | | | | | | | | | | | | | | | | | |
| Control/Nutrition | 313 | 0.95 | - | - | - | 235 | 0.93 | - | - | - | - | 311 | 1.59 | - | - | - | 233 | 1.35 | - | - | - | - |
| WSH/ WSH+Nutrition | 301 | 0.56 | 0.59 | 0.51 | 0.69 | 86 | 0.64 | 0.49 | 0.41 | 0.58 | 0.086 | 300 | 0.73 | -0.86 | -1.09 | -0.63 | 202 | 0.478 | -0.93 | -1.16 | -0.71 | 0.658 |
| Water | 165 | 0.61 | 0.64 | 0.54 | 0.77 | 120 | 0.93 | 0.70 | 0.57 | 0.87 | 0.519 | 165 | 0.85 | -0.72 | -1.02 | -0.43 | 86 | 0.795 | -0.51 | -0.78 | -0.24 | 0.301 |
| Handwashing | 155 | 0.96 | 1.01 | 0.96 | 1.07 | 202 | 0.47 | 1.01 | 0.94 | 1.09 | 0.998 | 154 | 1.56 | -0.06 | -0.26 | 0.15 | 120 | 1.316 | -0.03 | -0.30 | 0.24 | 0.859 |
| **Child Hands** | | | | | | | | | | | | | | | | | | | | | |
| Control/Nutrition | 295 | 0.93 | - | - | - | 220 | 0.89 | - | - | - | - | 285 | 1.88 | - | - | - | 218 | 1.64 | - | - | - | - |
| WSH/ WSH+Nutrition | 321 | 0.93 | 1.00 | 0.95 | 1.05 | 197 | 0.87 | 0.99 | 0.91 | 1.08 | 0.872 | 317 | 1.93 | 0.05 | -0.13 | 0.22 | 196 | 1.562 | -0.04 | -0.34 | 0.27 | 0.639 |
| **Sentinel Toys** | | | | | | | | | | | | | | | | | | | | | |
| Control/Nutrition | 327 | 0.77 | - | - | - | 360 | 0.81 | - | - | - | - | 321 | 0.72 | - | - | - | 357 | 0.60 | - | - | - | - |
| WSH/ WSH+Nutrition | 322 | 0.79 | 1.01 | 0.94 | 1.08 | 306 | 0.78 | 0.95 | 0.88 | 1.04 | 0.347 | 319 | 0.79 | 0.05 | -0.10 | 0.19 | 303 | 0.69 | 0.07 | -0.15 | 0.29 | 0.868 |
| **Flies Caught in Food Preparation Area** | | | | | | | | | | | | | | | | | | | | | |
| Control/Nutrition | 304 | 0.69 | - | - | - | 231 | 0.63 | - | - | - | - | 314 | 4.43 | - | - | - | 248 | 3.00 | - | - | - | - |
| WSH/ WSH+Nutrition | 345 | 0.59 | 0.95 | 0.85 | 1.06 | 233 | 0.60 | 1.03 | 0.89 | 1.19 | 0.364 | 356 | 3.86 | -0.83 | -2.18 | 0.52 | 234 | 3.39 | 0.46 | -0.72 | 1.64 | 0.169 |
| Sanitation | 203 | 0.68 | 0.96 | 0.79 | 1.18 | 181 | 0.69 | 0.91 | 0.71 | 1.18 | 0.759 | 207 | 3.58 | -1.08 | -2.51 | 0.35 | 188 | 2.99 | 0.52 | -1.13 | 2.18 | 0.162 |
| **Flies Caught in Latrine Area** | | | | | | | | | | | | | | | | | | | | | |
| Control/Nutrition | 304 | 0.69 | - | - | - | 231 | 0.63 | - | - | - | - | 304 | 3.90 | - | - | - | 231 | 3.04 | - | - | - | - |
| WSH/ WSH+Nutrition | 345 | 0.59 | 0.84 | 0.74 | 0.95 | 233 | 0.60 | 0.90 | 0.75 | 1.07 | 0.538 | 345 | 2.92 | -0.96 | -1.78 | -0.15 | 233 | 2.74 | -0.16 | -1.10 | 0.78 | 0.250 |
| Sanitation | 203 | 0.68 | 0.94 | 0.78 | 1.12 | 181 | 0.69 | 1.02 | 0.82 | 1.28 | 0.542 | 203 | 3.62 | -0.55 | -1.59 | 0.49 | 181 | 2.83 | 0.36 | -0.90 | 1.63 | 0.304 |

| | Two-Year Assessment | | | | | | | | | | | | | | | | | | | | | |
|---|---|---|---|---|---|---|---|---|---|---|---|---|---|---|---|---|---|---|---|---|---|---|
| | Prevalence | | | | | | | | | | | Concentration | | | | | | | | | | |
| | Wet Season | | | | | Dry Season | | | | | | Wet Season | | | | | Dry Season | | | | | |
| **Arms** | **N** | **Prev.** | **Prev. Ratio** | **95% Confidence Intervals** | | **N** | **Prev.** | **Prev. Ratio** | **95% Confidence Intervals** | | **Inter-action P-Value** | **N** | **Log10 Mean (CFU/ 100 mL)/ Mean (Flies)** | **Log Diff./ Fly Count** | **95% Confidence Intervals** | | **N** | **Log10 Mean (CFU/ 100 mL)/ Mean (Flies)** | **Log Diff./ Fly Count** | **95% Confidence Intervals** | | **Inter-action P-Value** |
| **Stored Water** | | | | | | | | | | | | | | | | | | | | | | |
| Control/Nutrition | 424 | 0.94 | - | - | - | 402 | 0.95 | - | - | - | - | 420 | 1.46 | - | - | - | 402 | 1.50 | - | - | - | - |
| WSH/ WSH+Nutrition | 364 | 0.73 | 0.78 | 0.71 | 0.85 | 188 | 0.76 | 0.84 | 0.79 | 0.89 | 0.199 | 364 | 1.14 | -0.32 | -0.47 | -0.17 | 362 | 1.238 | -0.26 | -0.39 | -0.12 | 0.515 |
| Water | 196 | 0.68 | 0.72 | 0.65 | 0.80 | 184 | 0.91 | 0.81 | 0.74 | 0.88 | 0.099 | 196 | 1.04 | -0.46 | -0.62 | -0.31 | 188 | 1.10 | -0.39 | -0.55 | -0.23 | 0.530 |
| Handwashing | 208 | 0.90 | 0.95 | 0.91 | 1.00 | 363 | 0.80 | 0.95 | 0.91 | 0.99 | 0.913 | 208 | 1.40 | -0.07 | -0.25 | 0.112 | 184 | 1.446 | -0.08 | -0.23 | 0.08 | 0.968 |
| **Child Hands** | | | | | | | | | | | | | | | | | | | | | | |
| Control/Nutrition | 448 | 0.86 | - | - | - | 412 | 0.94 | - | - | - | - | 439 | 1.54 | - | - | - | 405 | 1.90 | - | - | - | - |
| WSH/ WSH+Nutrition | 393 | 0.87 | 0.99 | 0.94 | 1.05 | 400 | 0.97 | 1.02 | 0.99 | 1.05 | 0.356 | 388 | 1.53 | -0.05 | -0.21 | 0.122 | 397 | 2.075 | 0.19 | 0.02 | 0.37 | 0.059 |
| **Sentinel Toys** | | | | | | | | | | | | | | | | | | | | | | |
| Control/Nutrition | 300 | 0.66 | - | - | - | 289 | 0.66 | - | - | - | - | 295 | 0.46 | - | - | - | 287 | 0.53 | - | - | - | - |
| WSH/ WSH+Nutrition | 269 | 0.66 | 0.97 | 0.87 | 1.08 | 249 | 0.62 | 0.95 | 0.84 | 1.08 | 0.874 | 268 | 0.52 | 0.006 | -0.14 | 0.149 | 249 | 0.43 | -0.09 | -0.25 | 0.07 | 0.381 |
| **Flies Caught in Food Preparation Area** | | | | | | | | | | | | | | | | | | | | | | |
| Control/Nutrition | 632 | 0.63 | - | - | - | 688 | 0.57 | - | - | - | - | 632 | 3.45 | - | - | - | 688 | 2.93 | - | - | - | - |
| WSH/ WSH+Nutrition | 389 | 0.60 | 0.98 | 0.88 | 1.09 | 407 | 0.61 | 1.04 | 0.93 | 1.17 | 0.405 | 389 | 2.89 | -0.60 | -1.49 | 0.30 | 407 | 3.01 | 0.17 | -0.41 | 0.74 | 0.155 |
| Sanitation | 320 | 0.68 | 1.04 | 0.94 | 1.14 | 286 | 0.59 | 0.97 | 0.85 | 1.11 | 0.462 | 320 | 3.66 | -0.26 | -1.39 | 0.88 | 286 | 2.71 | -0.22 | -0.82 | 0.39 | 0.946 |
| **Flies Caught in Latrine Area** | | | | | | | | | | | | | | | | | | | | | | |
| Control/Nutrition | 608 | 0.71 | - | - | - | 650 | 0.63 | - | - | - | - | 608 | 4.30 | - | - | - | 650 | 3.33 | - | - | - | - |
| WSH/ WSH+Nutrition | 386 | 0.69 | 1.00 | 0.93 | 1.09 | 390 | 0.65 | 1.03 | 0.93 | 1.14 | 0.696 | 386 | 3.46 | -0.61 | -1.23 | 0.00 | 390 | 3.44 | -0.10 | -0.73 | 0.53 | 0.256 |
| Sanitation | 312 | 0.72 | 1.01 | 0.91 | 1.13 | 275 | 0.69 | 1.08 | 0.95 | 1.23 | 0.435 | 312 | 3.82 | -0.83 | -1.84 | 0.17 | 275 | 3.55 | -0.08 | -0.72 | 0.57 | 0.201 |
